# DFT and Raman study of all-*trans* astaxanthin optical isomers

**DOI:** 10.1101/2021.03.11.434912

**Authors:** Guohua Yao, Shuju Guo, Wenjie Yu, Muhammad Muhammad, Jianguo Liu, Qing Huang

**Author notes:** These authors contributed equally to this work (Guohua Yao, Shuju Guo). Corresponding authors: Prof. Qing Huang, Address: CAS Key Laboratory of High Magnetic Field and Ion Beam Physical Biology, Institute of Intelligent Machines, Hefei Institutes of Physical Science, Chinese Academy of Sciences, Hefei 230031, China, Prof. Jianguo Liu, Address: CAS and Shandong Province Key Laboratory of Experimental Marine Biology, Center for Ocean Mega-Science, Institute of Oceanology, Chinese Academy of Sciences, Qingdao 266071, China.

## Abstract

Astaxanthin (AST) is a xanthophyll carotenoid widely distributed in aquatic animals, which has many physiological functions such as antioxidant, anti-inflammatory, anti-hypertensive and anti-diabetic activities. Astaxanthin has three optical isomers, including a pair of enantiomers (3*S*,3 ‘*S* and 3*R*,3 ‘*R*) and a meso form (3*R*,3 ‘*S*). Different optical isomers have differences in a variety of physiological functions. Traditionally, High Performance Liquid Chromatography (HPLC) can be used to distinguish these isomers. In this work, it’s found that Raman spectroscopy can be employed to distinguish the three optical isomers. Because the intensities of two Raman bands at 1190 cm^-1^ and 1215 cm^-1^ of three isomers are different. DFT calculations are performed and used to analyze the spectral differences. The calculation results show that the structures of these chiral isomers are not strictly mirror-symmetrical to each other, which leads to the difference in their Raman spectra.

**Highlights:**

- Raman spectroscopy can be utilized to distinguish three optical isomers of all-*trans* astaxanthin.
- The DFT-calculated spectrum is used to explain why the Raman bands of optical isomers at 1190 and 1215 cm^-1^ are different.
- The structural parameters of the three optical isomers have been identified.

## 1. Introduction

Astaxanthin (AST) is an orange-red, fat-soluble pigment, widely distributed in nature, particularly in aquatic animals including shrimp, crab and salmon. Astaxanthin, being a xanthophyll class of carotenoid, consists of two oxygenated ionone ring systems linked by a long conjugated double bond system.^1^ This part of astaxanthin is similar to β-carotene, lutein and zeaxanthin, and therefore they have some similar metabolic and physiological activities. But astaxanthin has a hydroxyl group and a keto group on each ionone ring, which makes it also have some special activities.^2^ In general, astaxanthin as an excellent antioxidant can scavenge free radicals in the body effectively, and it has a lot of physiological activities such as anti-inflammatory, anti-hypertensive, anti-diabetic, inhibiting tumorigenesis, lipid-lowering, cardiovascular disease prevention, and enhancing immunity functions. Therefore, it has attracted a lot of attention, recently in both researches and applications.^1, 3–4^ Astaxanthin products have been commercially applied in the human health nutrition, pharmaceutical/dietary supplements, and feed for aquatic animals.^4^

Astaxanthin has many kinds of geometrical isomers, since many conjugated double bonds in the polyene chain allow astaxanthin to exist as *cis-trans* isomers, with all-*trans* isomer predominant, some 9-*cis* and 13-*cis* isomers, very few 15-*cis* and di-*cis* isomers. ^5–6^ Each astaxanthin *cis-trans* isomer also has three optical isomers, including a pair of enantiomers (3*S*,3 ‘*S* and 3*R*,3 ‘*R*) and a meso form (3*R*,3 ‘*S*) due to the existence of two chiral carbons at the 3,3’ position.^7^ Astaxanthin from different sources have differences in the composition of optical isomers.^3^ The 3*S*,3 ‘*S* isomer is predominant form found in the microalga *Haematococcus pluvialis*,^8–9^ whereas the 3*R*,3 ‘*R* isomer is the main form in red yeast, *Phaffia rhodozyma*.^10^ Synthetic astaxanthin has a mixture of the three optical isomers (3*S*,3 ‘*S*, 3*R*,3 ‘*S* and 3*R*,3 ‘*R*) in a 1:2:1 ratio.^11^

The chiral structure of a biomolecule is usually closely related to its biofunction. Researchers are conducting various experiments to compare the bioactivities and functional properties of these isomers. Many studies have shown that 3*S*,3 ‘*S* astaxanthin exhibits higher antioxidant and anti-aging activities than 3*R*,3 ‘*R* and 3*R*,3 ‘*S* astaxanthin, both *in vivo* and *in vitro*, and indicated that different stereoisomers may have different targets in antioxidant and anti-aging mechanisms.^3, 12^ Specifically, these studies found that the levels of reactive oxygen species (ROS) in the worms fed with 3*S*,3 ‘*S*, 3*R*,3 ‘*R* and 3*R*,3 ‘*S* astaxanthin were reduced by 40%, 30%, 20% respectively. The 3*S*,3 ‘*S* isomer is predominant form found in the microalga, whereas the 3*R*,3’ *R* isomer is the main form in red yeast. Different tissues may have different preference for optical isomers of astaxanthin.^13^

Therefore, it is critical to identify the structures of different optical isomers. At present, there are very few technologies to distinguish there optical isomers. High Performance Liquid Chromatography (HPLC) is usually applied to separate and distinguish the optical isomers. In recent years, Raman spectroscopy technology has also been introduced to evaluate astaxanthin.^14–19^ However, to the best of our knowledge, Raman spectroscopy had not been used to study the optical isomers of astaxanthin. Therefore, in the present work, we introduced our method based on quantum chemistry and Raman spectroscopy to analyze the 3*S*,3 *S*, 3*R*,3 *S* and 3*R*,3 ‘*R* isomers of all-*trans* astaxanthin, systematically.

## 2. Experiment section

The all-*trans* isomer of astaxanthin was purchased from SenBeiJia Biological Technology Co., Ltd. Chiral isomers of all-*trans* astaxanthin were prepared from the all-*trans* astaxanthin on a Hanbon NP700 semipreparative HPLC system (Hanbon Sci. & Tech., Jiangsu, China) equipped with a CHIRALPAK IC column (5 μm, 250 mm × 10 mm, Daicel Corporation, Japan). The binary mobile phase consisted of A: methyl tert-butyl ether (MTBE) and B: acetonitrile. The solvent gradient was as follows: 0-24 min, 55% B. Peaks were detected at 470 nm by an UV-vis detector. The flow rate was set at 3.0 mL/min. Under these conditions, the chiral isomers fractions at the retention times of 15.622 min, 17.878 min, 20.534 min were collected and dried under a nitrogen stream to give the 3*S*,3 ‘*S*-, 3*R*,3 ‘*S*- and 3*R*,3 ‘*R*-all-*trans* astaxanthin separately, as shown in Figure 1.

**Figure 1.**
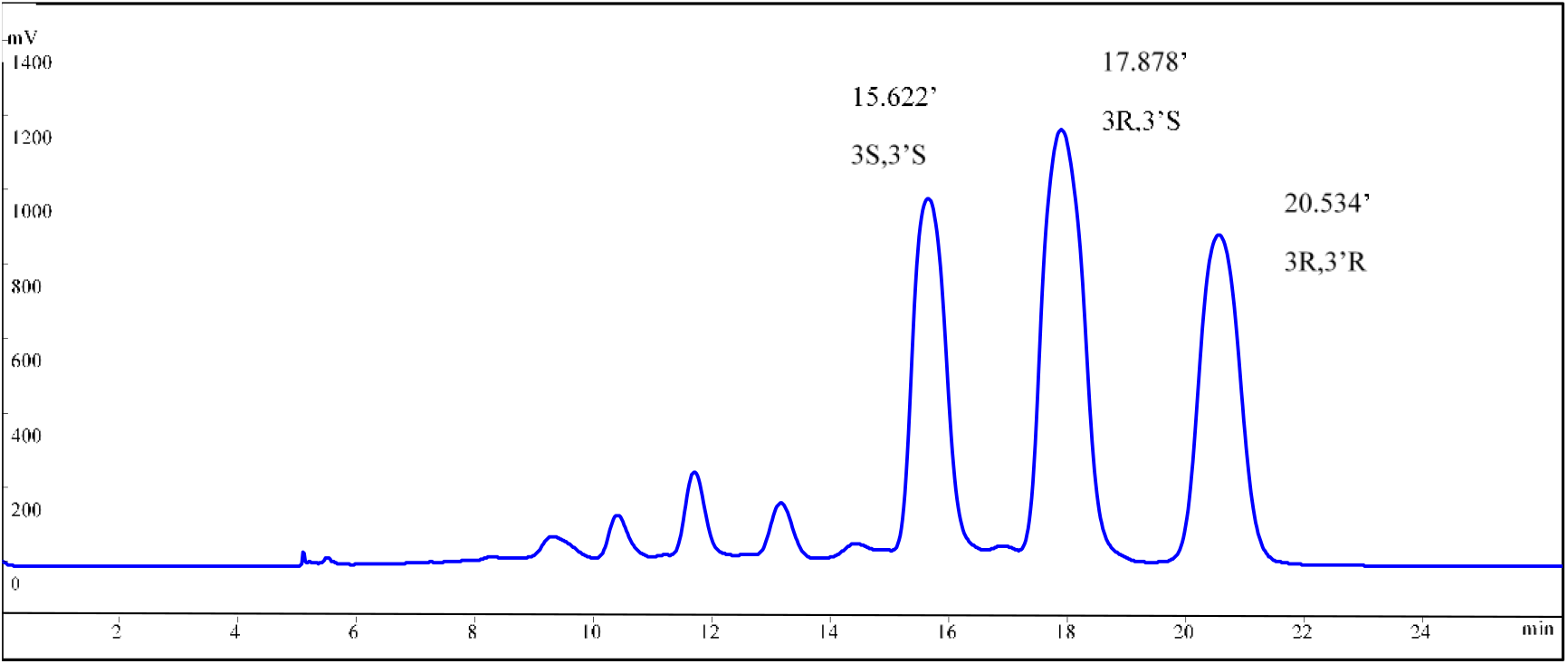
HPLC chromatograms of chiral isomers of all-*trans* astaxanthin

For the Raman spectral recording, the AST-sample powders are placed on a quartz plate for the Raman detections. All the Raman spectra were recorded in the 200-3700 cm^-1^ region using XploRA Raman spectrometer. (HORIBA JOBIN YVON) The laser power at the sample was ca. 1.2 mW, and the exposure time was 5 s. The spectra were measured with at least three repeats, then they were averaged, baseline corrected, and normalized.

## 3. Computational Details

Density functional theory calculations were carried out using Gaussian 09 software.^20^ All calculations were performed by applying the hybrid of Becke’s nonlocal three parameter exchange and correlation functional and the Lee-Yang-Parr correlation functional (B3LYP). The triple-zeta 6-311+G(d,p) split valence-shell basis set augmented by d polarization functions on heavy atoms and p polarization functions on hydrogen atoms as well as diffuse functions for heavy atoms was used.^21^ The geometries were fully optimized without any constraint on the planarity and the optimized geometries have no imaginary frequencies. The optimized structures and atom labels of three all-*trans* astaxanthin optical isomers are shown in Figure 2. Then the Raman spectra are simulated with a resolution of 4 cm^-1^ in the same level of theories. The vibrational frequency scaling factor of B3LYP/6–311+G (d,p) is 0.9808.^22^ Then the calculated Raman activities were converted into Raman intensities using the following relationship derived from the basic theory of Raman scattering: *I_i_* = *f*(*v*_0_ – *v_i_*)^4^*A_i_*/*v_i_* [1 – *exp*(−*hcV_i_*/*kT*)], where *v*_0_ is the exciting frequency (in unit of cm^-1^), *v_i_* is the vibrational frequency (in unit of cm^-1^) of the *i*th normal mode, *h, c*, and *k* are fundamental constants, and *f* is a suitably chosen common normalization factor for all peak intensities. Vibrational frequency assignments were made based on the results of the Gaussview program 5.0.8 version,^23^ and the potential energy distribution (PED) matrix was expressed in terms of a combination of local symmetry and internal coordinates.

**Figure 2.**
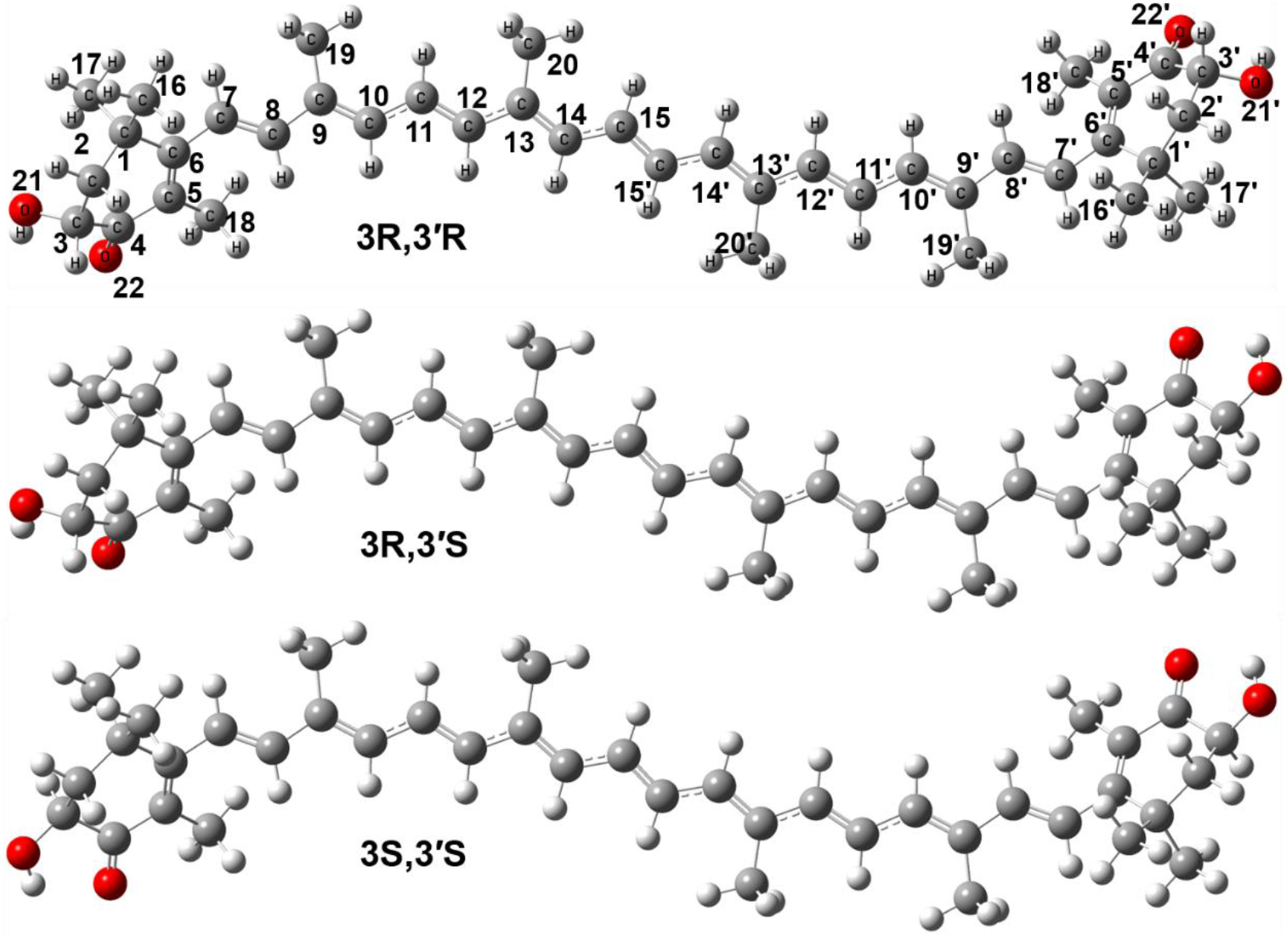
The optimized geometry structure and atom labeling of 3*R*,3 ‘*R*, 3*R*,3 ‘*S* and 3*S*,3 ‘*S*-all-*trans* astaxanthin.

## 4. Results and discussion

### 4.1 Experimental Raman spectra of the optical isomers

The experimental spectra of the 3*S*,3 ‘*S*-, 3*R*,3 ‘*S*- and 3*R*,3 ‘*R*-all-*trans* astaxanthin isomers are show in Figure 3. Obviously, they have similar Raman spectra. However, there are two weak bands near 1200 cm^-1^, and the intensities of the two bands of the three isomers are not the same. Therefore, we performed the split peak fitting for the bands in the range of 1100 cm^-1^ to 1240 cm^-1^, as shown in Figure 4. These two weak bands are at about 1190 and 1215 cm^-1^, respectively, their wavenumbers and normalized intensities are shown in the Table 1 and Figure 5. For these two bands, the wavenumbers of the three isomers are relatively similar, but the intensities are quite different. For the 1190 cm^-1^ band, the 3*R*,3 ‘*R* isomer has the lowest intensity, while the 3*S*,3 *S* isomer has the strongest intensity, while the intensity of 3*R*,3 ‘*S* isomer is between the two isomers, roughly the average of these two. For the 1215 cm^-1^, the 3*R*,3 ‘*R* isomer has the strongest intensity, and the 3*S*,3 ‘*S* isomer has the lowest intensity. The intensity ratio of 1215 cm^-1^ band to 1190 cm^-1^ band of 3*R*,3 ‘*R* isomer is 0.66, and the intensity ratio of 3*R*,3 ‘*S* is 0.48, and the ratio of 3*S*,3 ‘*S* is 0.24. Therefore, we can distinguish these three isomers based on the relative intensities of these two bands.

**Figure 3.**
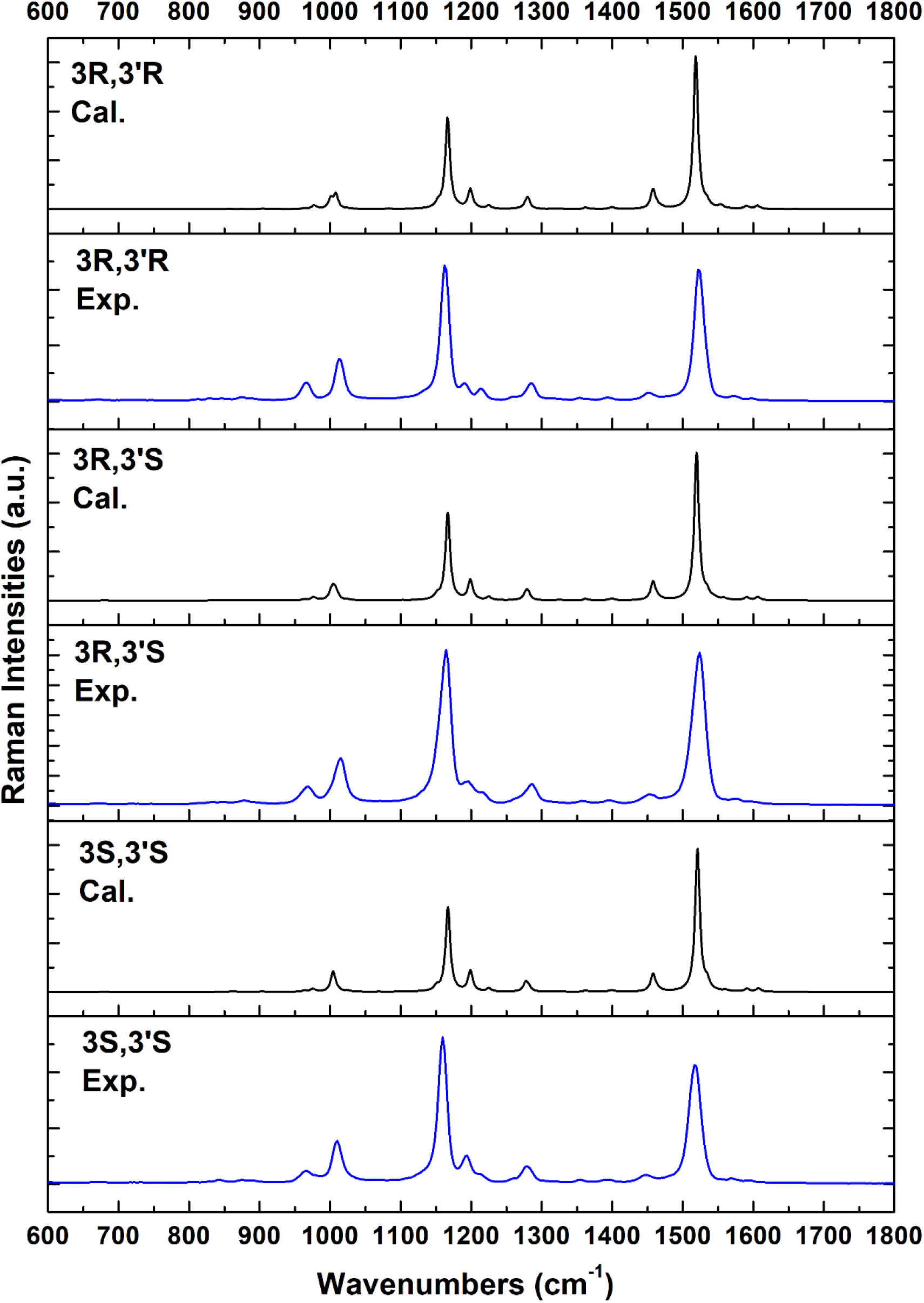
The experimental and calculated Raman spectra of 3*R*,3 ‘*R*, 3*R*,3 ‘*S* and 3*S*,3 ‘*S*-all-*trans* astaxanthin, respectively. The blue lines are the experimental Raman spectra, and the black lines are the simulated Raman spectra.

**Figure 4.**
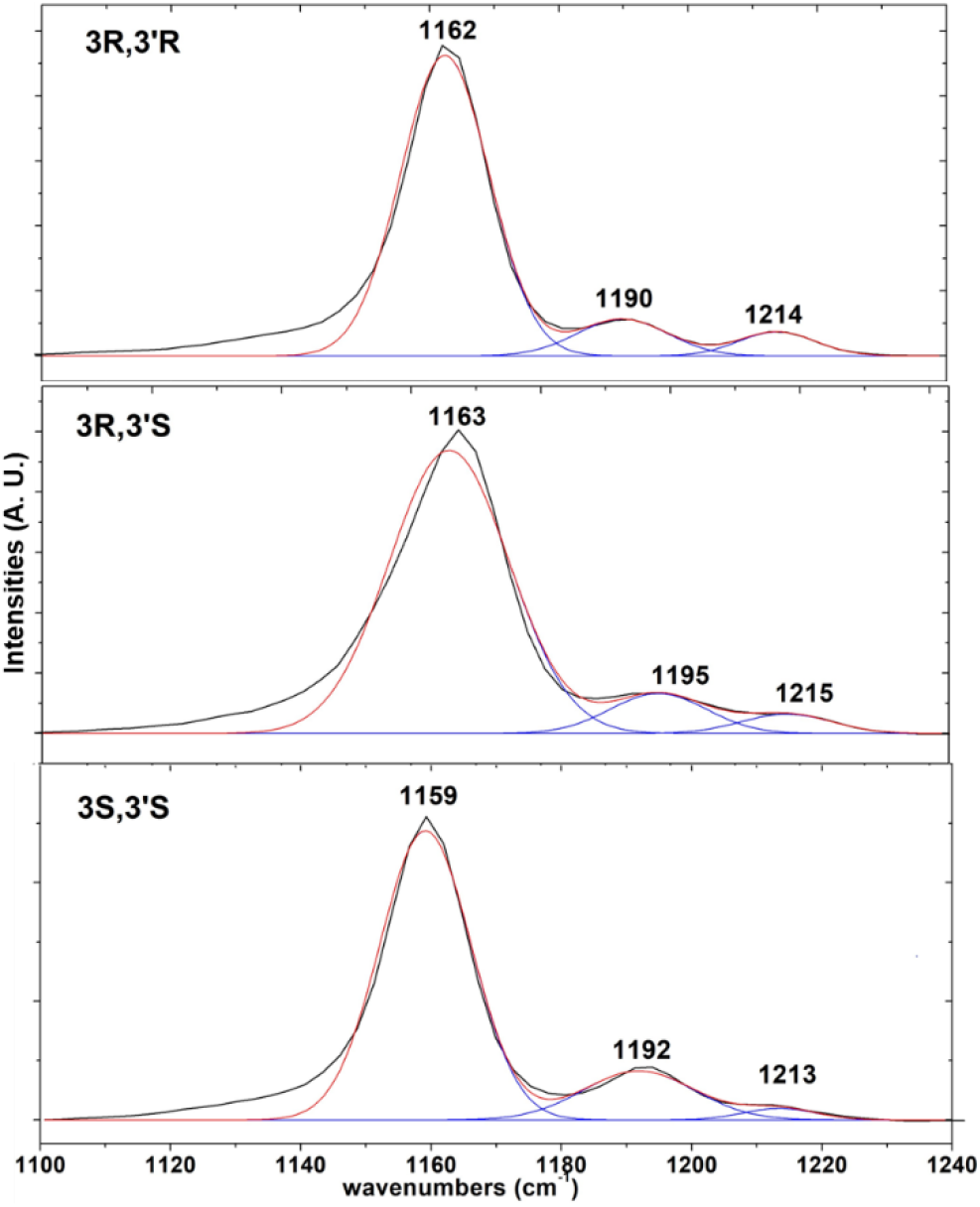
The multiple Raman peaks fitting of 3*R*,3 ‘*R*, 3*R*,3 ‘*S* and 3*S*,3 ‘*S*-all-*trans* astaxanthin between 1100 to 1240 cm^-1^.

**Table 1.** The experimental Raman wavenumbers and Raman intensities (a.u.), comparing with the calculated Raman bands and calculated Raman activity intensities (A4/AMU).

| isomer | exp. (cm<sup>-1</sup>) | exp. int. (a.u.) | cal. (cm<sup>-1</sup>) | cal. int. (A4/AMU) |
| --- | --- | --- | --- | --- |
| 3*R*,3 '*R* | 1190 | 0.117 | 1199 | 182022 |
|  | 1214 | 0.077 | 1225 | 31092 |
| 3*R*,3 '*S* | 1195 | 0.141 | 1199 | 189302 |
|  | 1215 | 0.068 | 1225 | 30498 |
| 3*S*,3 '*S* | 1192 | 0.169 | 1199 | 197449 |
|  | 1213 | 0.040 | 1225 | 29901 |

**Figure 5.**
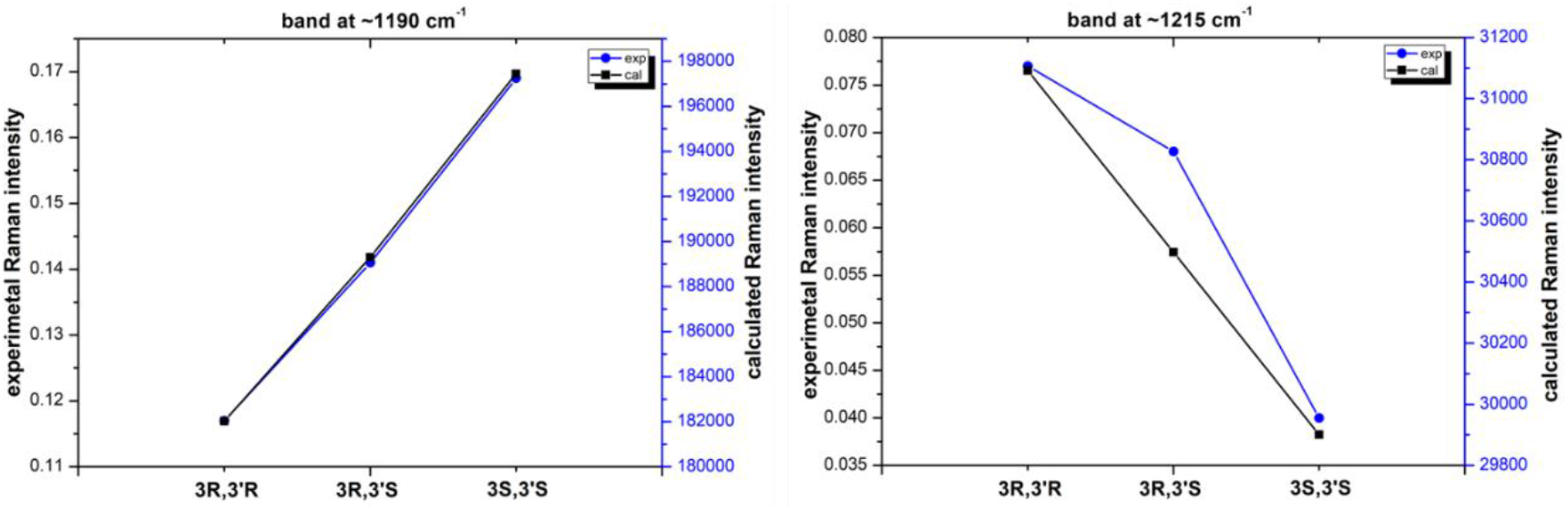
Comparison of the experimental and calculated Raman intensities of 3*R*,3 ‘*R*, 3*R*,3 ‘*S* and 3*S*,3 ‘*S*-all-*trans* astaxanthin. (left: band at about 1190 cm^-1^; right: band at about 1215 cm^-1^)

#### 4.2.1 Calculation of the structures and Raman spectra of the optical isomers

For most molecules, the Raman spectra of molecules with different chirality are the same. However, the Raman spectra of the three chiral astaxanthin are indeed different. The theoretical calculation work tries to analyze and explain this reason. Firstly, molecular structures of different chiral all-*trans* astaxanthin are constructed, and then their Raman spectra are calculated and compared with the experimental results, as shown in Figure 3, 4 and Table 1. Thereby, the structures of astaxanthin in accordance with the experimental results are achieved. Their optimization structures are shown in Figure 2, and the structural parameters are listed in Table 2.

**Table 2.**
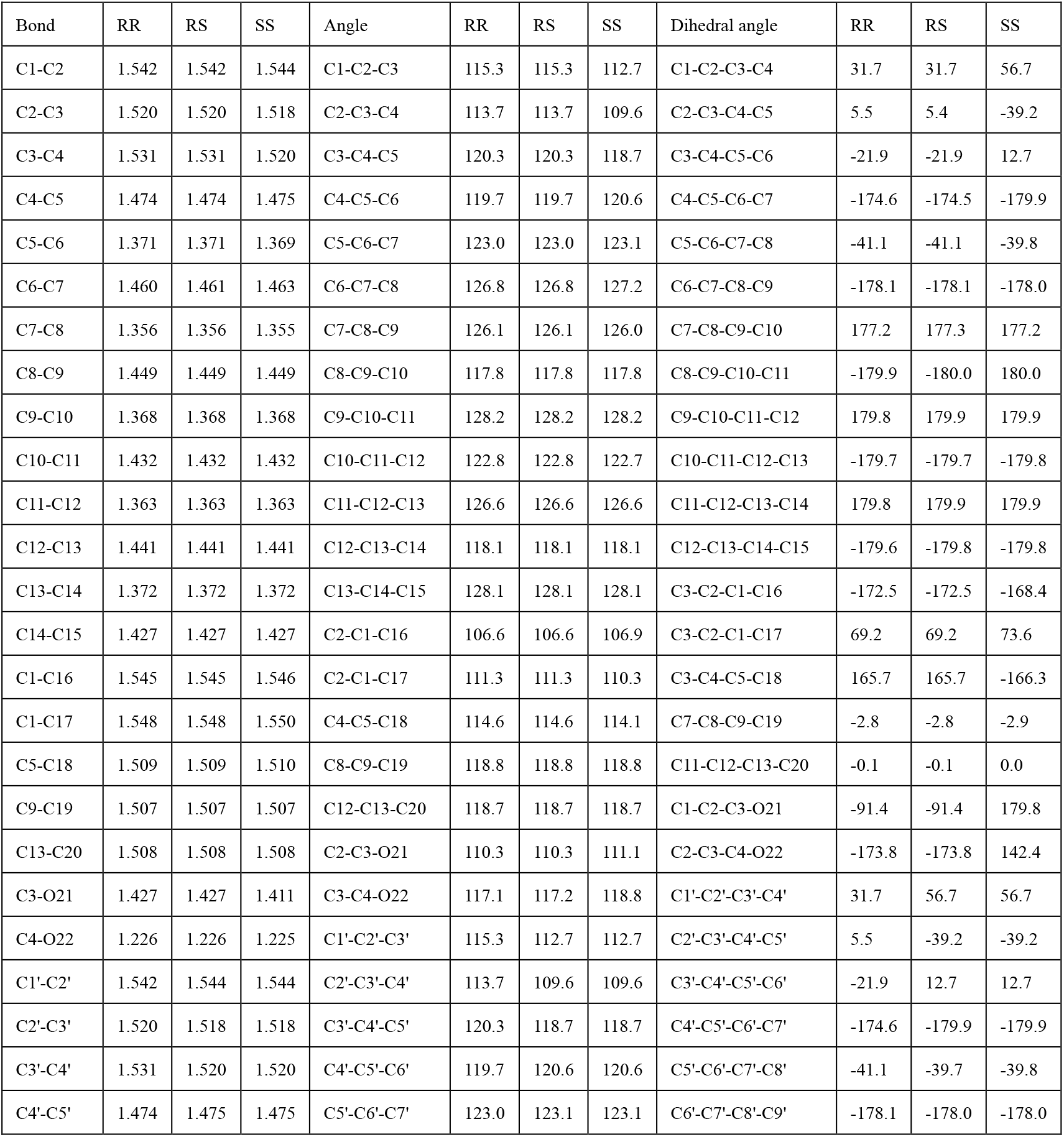
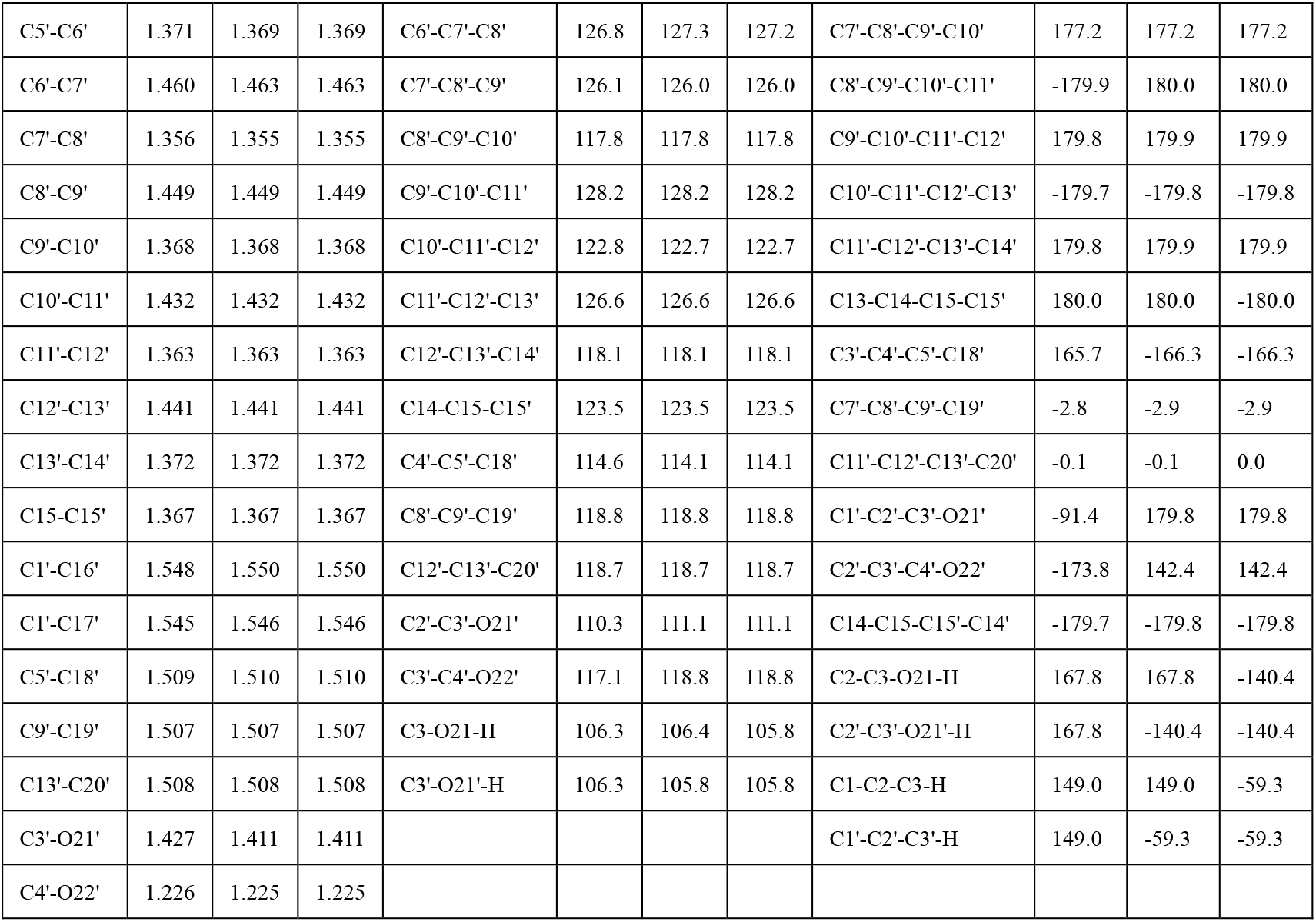
Bond lengths (Å), angles (°) and dihedral angles (°) of three optical isomers of all-*trans* astaxanthin calculated at B3LYP/6-311+G(d,p) Level.

Since C3 and C3’ are chiral carbon atoms in the astaxanthin molecule, in the *R* and *S* configurations, the configuration order of the groups attached to C3 is opposite. The C3 and C3’ of the 3*R*,3 ‘*R* isomer are all with the *R* chiral, C3 and C3’ of the 3*S*,3 ‘*S* isomer are all with the *S* chiral. The configurations of 3*R*,3 ‘*R* and 3*S*,3 ‘*S* are the enantiomers of each other. C3 of the 3*R*,3 ‘*S* isomer is *R* chiral and C3’ of the 3*R*,3 ‘*S* isomer is *S* chiral, therefore 3*R*,3 *S* isomer is a meso configuration.

The structure 3*R*,3 ‘,*R*-all-*trans* astaxanthin isomer has C2 symmetry, the structural parameters of the left half (C1 to C15) and the right half (C1’ to C15’) of the molecule are the same. For instance, the C3-O21-H and C3’-O21’-H angles are the all 106.3°, the C1-C2-C3-C4 and C1’-C2’-C3’-C4’ dihedral angles are all 31.7°. Besides, the structure 3*S*,3 *S*-all-*trans* astaxanthin isomer also has C2 symmetry. The C1-C2-C3-C4 and C1’-C2’-C3’-C4’ dihedral angles of 3*S*,3 ‘*S* are all 56.7°. But 3*R*,3 ‘*S* isomer is asymmetric (C1 point group). The structural parameters of left half of 3*R*,3 ‘*S* isomer is nearly the same with the 3*R*,3 ‘*R* isomer, while the structural parameters of right half of 3*R*,3 ‘*S* isomer is nearly the same with the 3*S*,3 ‘*S* isomer. For instance, the C1-C2-C3-C4 dihedral angle is 31.7°, C1’-C2’-C3’-C4’ dihedral angle is 56.7°.

The structural parameters between the three optical isomers are also compared. The bond length parameters of these three optical isomers are very close, while their angle parameters, especially dihedral angle parameters, are quite different. Their angle and dihedral angle parameters of the long conjugated double bond system are similar. The difference mainly lies on the oxygenated ionone ring systems. For example, C2-C3-C4-C5 angles of 3*R*,3 ‘*R* and 3*R*,3 ‘*S* isomers are 5.5° and 5.4°, respectively, while C2-C3-C4-C5 angle of 3*S*,3 ‘*S* is −39.2°. The C2-C3-O21-H of 3*R*,3 ‘*R* isomer is 167.8°, the C2’-C3’-O21’-H of 3*S*,3 ‘*S* isomer is −140.4°. Therefore, the structures of 3*R*,3 ‘*R* and 3*S*,3 ‘*S* are not mirror-symmetrical to each other, although their configurations are mirror-symmetrical to each other.

Since the structures of 3*R*,3 ‘*R* is not mirror-symmetrical to 3*S*,3 ‘*S*, the difference in structure may naturally lead to the difference in Raman spectra. The simulated Raman spectra of all-*trans* astaxanthin optical isomers are shown in Figure 3. It can be seen that in general, the peak intensity and wavenumber of the theoretical spectrum are quite consistent with that of the experimental spectrum. The wavenumbers, intensities and PEDs of the calculated Raman bands of three optical isomers are also listed in Table 3, 4, 5. The wavenumbers of the same experimental band of different isomers only have a difference of less than 8 wavenumbers. While the wavenumbers of the same simulated band of different isomers only have a difference of less than 4 wavenumbers. The assignments of different isomers are also similar, with only slight difference in the ratio of energy.

**Table 3.** Comparison between the experimental and theoretical calculated Raman spectra of the 3*R*,3 ‘*R*-all-*trans* astaxanthin, the assignments and PED of vibrational modes at B3LYP/6-311+G(d,p) Level.

| Exp | Int | Cal | Int | Assignments (PED %) |
| --- | --- | --- | --- | --- |
| 1594 | w | 1606 | 39972 | str C7=C8 (13), str C7'=C8' (13) |
| 1567 | w | 1590 | 34924 | str C9=C10 (19), str C9'=C10' (19) |
| 1516 | vs | 1518 | 1745714 | str C13=C14 (12), C13'=C14' (12), C15=C15' (10) |
| 1447 | w | 1458 | 138513 | sciss C20H3 (16), sciss C20'H3 (16) |
| 1388 | w | 1400 | 14334 | sciss C19H3 (11), sciss C19'H3 (11) |
| 1354 | w | 1362 | 10363 | rock C14-H (10), rock C14'-H (10) |
| 1279 | m | 1280 | 114167 | rock C11-H (16), rock C11'-H (16) |
| 1214 | w(0.077) | 1225 | 31092 | str C12-C13 (11), C12'-C13' (11), C10-C11 (10), C10'-C11' (10) |
| 1190 | m(0.117) | 1199 | 182022 | str C8-C9 (13), C8'-C9' (13) |
| 1162 | vs | 1167 | 851324 | str C14-C15 (15), C14'-C15' (15), C10-C11 (12), C10'-C11' (12) |
| 1010 | s | 1008 | 119881 | wag C20H3 (12), C20'H3 (12), C19H3 (11), C19'H3 (11) |
| 965 | m | 977 | 21776 | bend out C8'-H (20), C8-H (20), C15'-H (10), C15-H (10) |

Abbreviations: PED, potential energy distribution (PED above 10 percent are listed); above 10 percent. Exp, the wavenumber of experimental Raman shift (in $\text{cm}^{-1}$ ); Cal, the wavenumber of calculated Raman shift (in $\text{cm}^{-1}$ ); Int, intensity (the calculated Raman activity intensity is in $\text{A}^4/\text{AMU}$ ); $\Delta v$ , the calculated Raman shift minus experimental Raman shift of the vibrational mode (in $\text{cm}^{-1}$ ); s, strong; m, medium; w, weak; vw, very weak; sh, shoulder peak; bend, bending; bend out, out-of-plane bending; rock, rocking; sciss, scissoring; str, stretching; wag, wagging.

**Table 4.** Comparison between the experimental and theoretical calculated Raman spectrum of the 3*R*,3 ‘*S*-all-*trans* astaxanthin, the assignments and PED of vibrational modes at B3LYP/6-311+G(d,p) Level.

| Exp | Int | Cal | Int | Assignments (PED %) |
| --- | --- | --- | --- | --- |
| 1596 | w | 1607 | 39632 | str C7=C8 (15), str C7'=C8' (11) |
| 1575 | w | 1591 | 33712 | str C9'=C10' (19), str C9=C10 (18) |
| 1524 | vs | 1520 | 1693637 | C13'=C14' (12), str C13=C14 (11), C15=C15' (10) |
| 1452 | w | 1458 | 131779 | sciss C20'H3 (22), sciss C20H3 (15) |
| 1393 | w | 1400 | 12636 | bend C3'-O20'-H (10) |
| 1356 | w | 1362 | 13095 | rock C14'-H (17), rock C14-H (16) |
| 1287 | m | 1281 | 52428 | twist C2'H2 (19), bend C3'-O20'-H (10) |
| 1215 | w(0.068) | 1225 | 30498 | str C12'-C13' (12), C12-C13 (11), C10'-C11' (11), C10-C11 (10) |
| 1195 | m(0.141) | 1199 | 189302 | str C8-C9 (13), C8'-C9' (13) |
| 1163 | vs | 1167 | 815384 | str C14-C15 (15), C14'-C15' (15), C10-C11 (13), C10'-C11' (12) |
| 1015 | s | 1004 | 84514 | C20'H3 (13), C19'H3 (13) |
| 970 | m | 976 | 20439 | C8'-H (23) |

**Table 5.** Comparison between the experimental and theoretical calculated Raman spectrum of the 3*S*,3 ‘*S*-all-*trans* astaxanthin, the assignments and PED of vibrational modes at B3LYP/6-311+G(d,p) Level.

| Exp | Int | Cal | Int | Assignments (PED %) |
| --- | --- | --- | --- | --- |
| 1596 | w | 1607 | 39609 | str C7=C8 (13), str C7'=C8' (13) |
| 1572 | w | 1591 | 32857 | str C9=C10 (18), str C9'=C10' (18) |
| 1522 | vs | 1521 | 1648698 | str C13=C14 (11), C13'=C14' (11), C15=C15' (10) |
| 1452 | w | 1458 | 99254 | sciss C20H3 (30), sciss C20'H3 (30) |
| 1394 | w | 1400 | 10766 | bend C3-O20-H (15), bend C3'-O20'-H (15) |
| 1354 | w | 1362 | 15347 | rock C14-H (13), rock C14'-H (13) |
| 1285 | m | 1277 | 85817 | rock C11-H (14), rock C11'-H (14) |
| 1213 | w(0.040) | 1225 | 29901 | str C10-C11 (11), C10'-C11' (11), C12-C13 (10), C12'-C13' (10) |
| 1192 | w (0.169) | 1199 | 197449 | str C8-C9 (13), C8'-C9' (13) |
| 1159 | vs | 1167 | 781333 | str C14-C15 (15), C14'-C15' (15), C10-C11 (13), C10'-C11' (13) |
| 1013 | s | 1004 | 171782 | wag C20H3 (12), C20'H3 (12), C19H3 (11), C19'H3 (11) |
| 965 | m | 975 | 23980 | bend out C8-H (14), C8'-H (14) |

The structure of 3*R*,3 ‘*R* and 3*S*,3 ‘*S* all-*trans* astaxanthin molecules have C2 symmetry, therefore, the distributions of their vibration modes are also symmetric. And the distributions of 3*R*,3 ‘*S* is not symmetric due to its asymmetric structure. In the 1500-1700 cm^-1^ regions, the bands are mainly from the stretching vibrational modes of carbon-carbon double bonds. The most intense experimental band at about 1520 cm^-1^ is from the symmetry stretching of carbon-carbon double bonds of the whole polyene chain, the PEDs of C13=C14, C13-C14’ and C15=C15’ stretching vibrations are above 10 percent, and the PED of C11=C12, C11’=C12’, C9=C10 and C9=C10’ are less than 10 percent. In the 800 to 1500 cm^-1^ regions, the bands are mainly from the stretching vibration of carbon-carbon single bonds and bending of C-H bonds. The strong band at about 1010 cm^-1^ is from wagging vibration of methyl groups on the chain, which are C20H_3_, C20’H_3_, C19H_3_, C19’H_3_. The very strong band at 1160 cm^-1^ is mainly from symmetry stretching of C14-C15, C14’-C15’, C10-C11, C10’-C11’ bonds. The band at about 1190 cm^-1^ mainly is from the stretching of C8-C9, C8’-C9’ bonds. The weak band at about 1215 cm^-1^ is from the stretching of carbon-carbon single bonds of the polyene chain, mainly from C10-C11, C10’-C11’, C12-C13, C12’-C13’ bonds. As shown in table 1 and Figure 5, the simulated Raman activity intensities of 1190 cm^-1^ and 1215 cm^-1^ bands of three isomers agree with the experiment. For the 1190 cm^-1^ band, the 3*R*,3 ‘*R* isomer has the lowest Raman activity intensity, and the 3*S*,3 ‘*S* isomer has the strongest Raman activity intensity, while the Raman activity intensity of 3*R*,3 ‘*S* isomer is the average of these two. On the contrary, for the 1215 cm^-1^ band, the 3*R*,3 ‘*R* isomer has the strongest Raman activity intensity, the 3*S*,3 ‘*S* isomer has a lowest Raman activity intensity, while the Raman activity intensity of 3*R*,3 ‘*S* isomer has the average of these two.

Although these easily identifiable Raman bands are mainly due to the vibration of atoms in the conjugated long chain. But in most vibration modes, each atom is more or less involved in vibration. The structural difference in the oxygenated ionone ring also slightly affects the Raman wavenumbers and activity intensities. Due to the structural difference of 3*R*,3 ‘*R*, 3*R*,3 ‘*S* and 3*S*,3 ‘*S* isomers, the Raman activity intensities of their these two bands are correspondingly different. So that different chiral isomers can be distinguished in the experimental spectrum. This phenomenon also exists in other molecules. For example, the simplest amino acid Glycine, appears mainly in three crystals modifications: α, β and γ.^24–25^ Even the new formation like δ and ζ was observed under different physical conditions, and their Raman spectra would be different.^26^ Due to the different interactions between molecules in different forms, the molecular structure has changed, resulting in different wavenumbers or intensities of their Raman spectra.

## 5. Conclusion

In this work, the Raman spectra of three optical isomers of all-*trans* astaxanthin were studied, which are 3*R*,3 ‘*R*, 3*R*,3 ‘*S* and 3*S*,3 ‘*S* isomers. It was found that the Raman intensities of three isomers at about 1190 cm^-1^ and 1215 cm^-1^ are different. The intensity of the 1190 cm^-1^ band of 3*R*,3 ‘*R* isomer is lowest among these isomers, and the intensity of the 1215 cm^-1^ band is strongest. The intensity of the 1190 cm^-1^ band of 3*S*,3 ‘*S* isomer is strongest among these isomers, while the intensity of the 1215 cm^-1^ band is lowest. While two bands of 3*R*,3 ‘*S* isomer are of intermediate strength. According to the intensity ratio of these two bands, the three optical isomers can be distinguished. DFT calculations are performed to explain the reasons for spectral differences. With the structure of the isomers being optimized, their Raman spectra being simulated, and their vibration modes being analyzed, the calculation results show that the structures of these chiral isomers are not mirror-symmetrical to each other. The difference in structure leads to the difference in their Raman spectra. As such, this work provides the chiral recognition of astaxanthin molecules, and may promote the study of the physiological functions of chiral astaxanthin molecules.

## Acknowledgement

This work was supported by the National Natural Science Foundation of China (11635013, 11775272, 21904127, 11475217 and U1706209), the Anhui Provincial Natural Science Foundation (1908085MB53) and Laboratory for Marine Biology and Biotechnology, Qingdao National Laboratory for Marine Science and Technology (MS2019NO01).

